# Restructuring of olfactory representations in the fly brain around odor relationships in natural sources

**DOI:** 10.1101/2023.02.15.528627

**Authors:** Jie-Yoon Yang, Thomas F. O’Connell, Wei-Mien M. Hsu, Matthew S. Bauer, Kristina V. Dylla, Tatyana O. Sharpee, Elizabeth J. Hong

**Affiliations:** Division of Biology & Biological Engineering, California Institute of Technology, Pasadena, CA, USA; Computational Neurobiology Laboratory, Salk Institute for Biological Studies, La Jolla, CA, USA; Department of Physics, University of California, San Diego, La Jolla, CA, USA

## Abstract

A core challenge of olfactory neuroscience is to understand how neural representations of odor are generated and progressively transformed across different layers of the olfactory circuit into formats that support perception and behavior. The encoding of odor by odorant receptors in the input layer of the olfactory system reflects, at least in part, the chemical relationships between odor compounds. Neural representations of odor in higher order associative olfactory areas, generated by random feedforward networks, are expected to largely preserve these input odor relationships^1–3^. We evaluated these ideas by examining how odors are represented at different stages of processing in the olfactory circuit of the vinegar fly *D. melanogaster*. We found that representations of odor in the mushroom body (MB), a third-order associative olfactory area in the fly brain, are indeed structured and invariant across flies. However, the structure of MB representational space diverged significantly from what is expected in a randomly connected network. In addition, odor relationships encoded in the MB were better correlated with a metric of the similarity of their distribution across natural sources compared to their similarity with respect to chemical features, and the converse was true for odor relationships encoded in primary olfactory receptor neurons (ORNs). Comparison of odor coding at primary, secondary, and tertiary layers of the circuit revealed that odors were significantly regrouped with respect to their representational similarity across successive stages of olfactory processing, with the largest changes occurring in the MB. The non-linear reorganization of odor relationships in the MB indicates that unappreciated structure exists in the fly olfactory circuit, and this structure may facilitate the generalization of odors with respect to their co-occurence in natural sources.

## INTRODUCTION

The search for organizing principles of olfaction has often focused on relating the chemical structure or physicochemical properties of odorants to their percept^4^. This approach is principled since odors are detected by their molecular interactions with large families of structurally diverse odorant receptor (OR) proteins expressed in ORNs^5^. Recently, significant inroads have been made in predicting a molecule’s odor from its structure^6,7^ but developing a generalized relationship between odorant structure and perception across the space of all possible odor stimuli remains challenging because of discontinuities in this relationship: small changes in structure often result in dramatic changes in a molecule’s odor^8,9^. This gap in understanding motivates a search for additional organizational axes of odor space to complement structure-based approaches towards gaining a better understanding of what determines a molecule’s smell.

Another important property of odorants is how they are organized relative to one another in natural environments. Odors from natural sources are typically complex mixtures of dozens to hundreds of monomolecular odorants, the composition of which is controlled by the conserved biochemical and metabolic processes in the source^10,11^. The relative abundance or ratios of volatiles in natural odor profiles can provide information about the value or state of the odor source^12–13^, for instance, if microbes that promote fermentation or spoilage are dominant. Thus, the odor space of the natural world is highly structured, and this structure often contains information about the identity or ethological value of the odor source.

We investigated how representations of odor at different stages of processing in the brain of the vinegar fly *Drosophila melanogaster* relate to different odor properties, focusing on their chemical properties or their relative abundances in behaviorally significant natural odor sources like food. The fly has a compact olfactory system with a similar overall circuit architecture to its vertebrate counterpart^14^. All ORNs that express the same OR project to a common synaptic compartment, called a glomerulus, in the antennal lobe (AL), and the dendrites of uniglomerular second-order projections neurons (PNs) extend into a single glomerulus^15^. Thus, each glomerulus, corresponding to a specific OR, represents a fundamental unit of olfactory processing. A major target of PN output from the AL is the mushroom body (MB), a cerebellum-like associative center in the fly brain that encodes representations of odor identity^16^. Wiring of PN inputs to Kenyon cells (KCs), the principal neurons of the MB, is probabilistic: each of the ~2000 KCs integrates input from a subset of PNs comprising ~10% of the ~50 olfactory glomeruli in the system^17–19^. KCs have high spiking thresholds and act as coincidence detectors that fire only when multiple input PNs are co-active^20,21^, and local feedback inhibition between KCs is provided from the arborizations of an unusual single, large GABAergic neuron called the APL^22^.

This circuit architecture recodes dense, distributed representations of odor in the ~50 glomeruli of the PN layer into a sparse, high-dimensional representation in the MB layer^3,23–25^ that facilitates pattern separation and linear decoding by a smaller number of MB output neurons. Theoretical studies of cerebellum-like circuits, characterized by expansion (PN input onto KCs) and reconvergence (KC output onto MB output neurons), emphasize the role of random, unstructured input for decorrelating activity patterns and maximizing the dimensionality of representations^26,27^. Such features would promote efficient memory storage and reduced synaptic inference during stimulus-specific associative learning and recall.

Feedforward random network models of the MB predict that odor relationships encoded in the MB should be strongly decorrelated, invariant across individual brains, and should preserve stimulus relationships encoded at the level of ORN input ^1,28–30^. However, recent large-scale EM reconstructions of MB synaptic connectivity demonstrated that some PN inputs are structured.

In particular, inputs from glomeruli tuned to odors common in food are more likely to converge onto the same KC targets^19^, though the functional impact on MB representations of odor remains to be determined. Whereas random networks maximize coding capacity and promote the separability of odor representations throughout odor space, structured networks can correlate specific odor representations to promote generalization between odors sharing particular ethological meaning. We investigated how odor coding in the fly olfactory circuit balances these competing needs.

## RESULTS

### Population imaging of odor representations in the MB at cellular resolution

In the arc of sensorimotor transformation, representations of odor identity encoded in KC activity patterns represent the output from the sensory arm of the pathway, which is flexibly coupled to distinct downstream outputs and behaviors^31^. Thus, we began by investigating representations of odor in the MB. The *Drosophila* olfactory circuit is the most comprehensively mapped metazoan olfactory system, with the tuning of approximately half of the odorant receptors to a large panel of 109 odors described by the Hallem dataset^32^. We selected 24 monomolecular odors that spanned OR input space (Figure 1A) and investigated the extent to which those relationships at the periphery are maintained in the MB.

**Figure 1:**
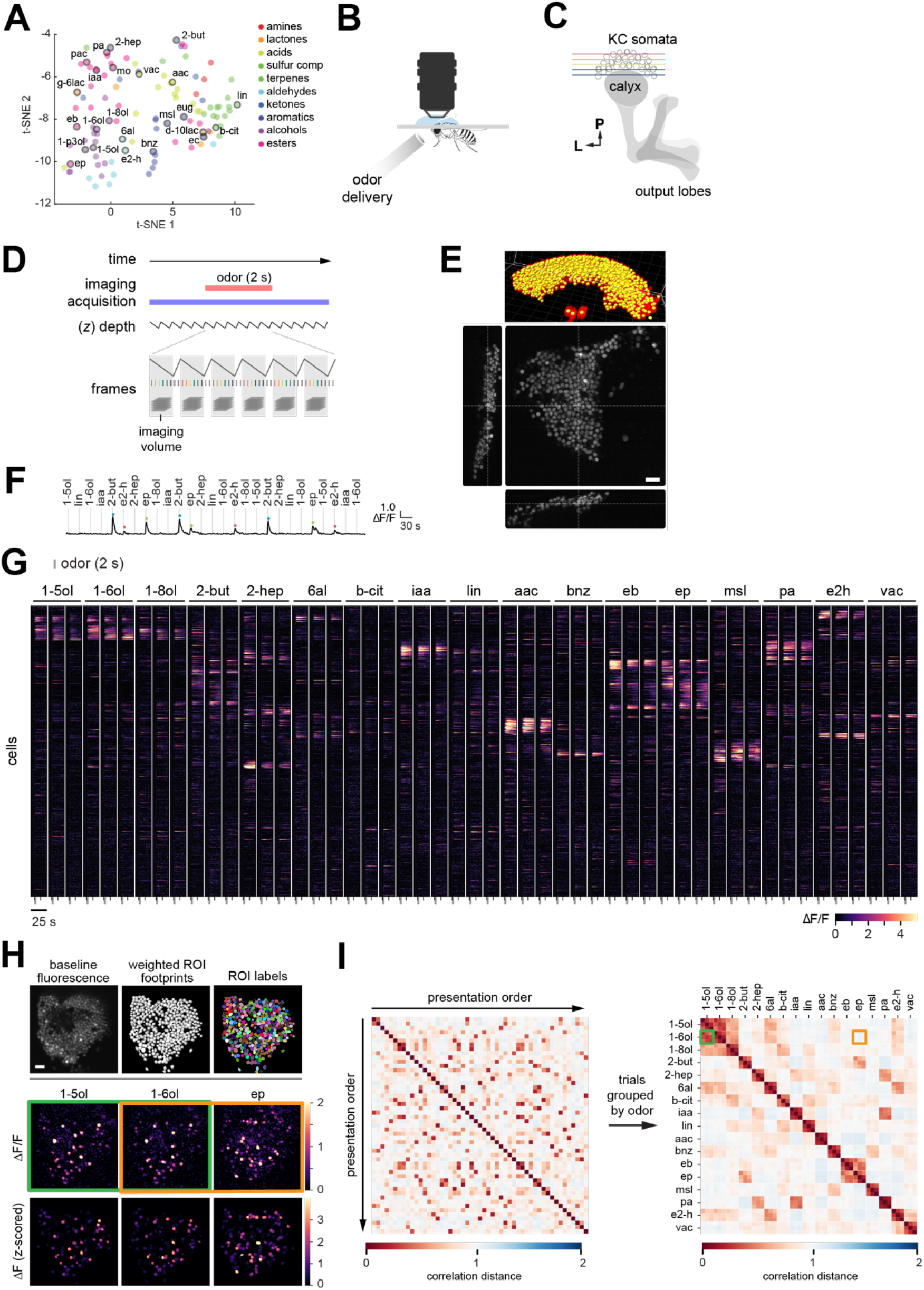
Population representations of odor in the fly MB at cellular resolution. **A)** bedding of 109 odors based on the patterns of activity they elicit across 24 fly ORs in the Hallem dataset. A subset (open grey symbols) spanning the odor space was selected for measurement in the MB. **B)** Odors were delivered to the antennae of immobilized flies expressing nuclear-localized GCaMP6s in all KCs, while imaging from KC somata with a two-photon microscope. **C)** Imaging volumes comprising ~11 planes through the KC layer capture the activity of >85% KCs in an MB. **D)** Configuration of volumetric imaging trial (3 Hz sweep rate). **E)** Reconstruction of 3D ROIs corresponding to each KC from interleaved high-resolution anatomical imaging stacks. Scale bar, 10 *μ*m. **F)** Example odor-evoked calcium signals in an imaging block from a single KC. **G)** Population representations in ~1800 KCs to 17 odors in the MB from a representative fly. Each row is a cell and each column is a trial. Cells are sorted by odor tuning. **H)** Top: baseline fluorescence (left), weighted ROI masks of KCs (middle), and ROI lablels corresponding to individual KCs (right) from a single imaging plane in an MB. Bottom: odor-evoked patterns of KC activity in response to 1-pentanol, 1-hexanol, and ethyl propionate. Scale bar, 10 *μ*m. I) Matrix of pairwise correlation between KC population responses in individual odor trials, where trials are shown in presentation order (left) or grouped by odor (right). KC responses elicited by 1-hexanol were more similar to those elicited by 1-pentanol (green) than by ethyl propionate (orange). Flies had genotype 20xUAS-nls-OpGCaMP6s-p10 (III)/+; OK107-Gal4 (IV)/+. Odors are pentyl acetate (pa), isoamyl acetate (iaa), methyl octanoate (mo), ethyl butyrate (eb), ethyl propionate (ep), 1-pentanol (1-5ol), 1-hexanol (1-6ol), 1-penten-3-ol (1-p3ol), 1-octanol (1-8ol), 2-heptanone (2-hep), 2-butanone (2-but), hexanal (6al), E2-hexenal (e2-h), benzaldehyde (bnz), methyl salicylate (msl), eugenol (eug), ethyl cinnamate (ec), linalool (lin), ß-citronellol (b-cit), acetic acid (aac), propionic acid (pac), valeric acid (vac), γ-hexalactone (g-6lac), δ-decalactone (d-10lac).

We used in vivo volumetric two-photon microscopy to image odor-evoked calcium signals in the MB of flies expressing nuclear-localized GCaMP6s selectively in KCs (directed by the *OK107-Gal4* driver^33^) (Figure 1B-C). In pilot experiments using cytoplasmic GCaMP6f as the calcium reporter, the small size (~2-3 μm) and tight, regular packing of KC somata (Figure 1E) presented challenges for motion correction, good cellular segmentation, and reliable pixel assignment to individual KCs over the course of an imaging session, with poor trial-to-trial reliability in odor panels larger than 8 odors. To expand the size of odor panels that could be evaluated in a single brain, we turned to measuring nuclear calcium, which has slower response dynamics, but strongly correlated response amplitudes compared to cytoplasmic calcium^34,35^. Localization of the calcium indicator to nuclei resulted in a several pixel gap between KCs that facilitated reliable cellular segmentation (Figure 1E, H) and enabled recording of the representations of between 8 to 17 odors in the same MB.

Flies were presented odors in pseudo-random sequences while rapidly z-scanning through the KC cell body layer. Following volumetric motion correction, odor-evoked KC signals were extracted using the Suite2P software package^36^. In brief, after correcting for motion in each plane, regions of interests (ROIs) representing each KC were extracted (Figure 1H). Although cell detection in Suite2P is usually neural activity-based, we extracted the spatial footprint of each cell by performing anatomical segmentation on time-averaged images, resulting in detection of between ~85-95% of the expected number of KCs. Since KC odor responses are sparse and many KCs do not respond to any odor in even a relatively large panel, this adjustment enabled reliable estimates of KC response rates. For a subset of experiments, odor responses were registered to KCs across different functional movies collected from the same MB by alignment of ROIs to 3D anatomical models of KC somata constructed from high-resolution structural images through the MB (Figure 1E; Figure S1A).

Reproducible odor-specific response dynamics were observed in some cells (Figure 1F), but, given the overall slow kinetics of nuclear-localized GCaMP6s, we focused our analysis in this study on the peak amplitude of odor-evoked responses. KC population responses were stable, odor-specific, and reliable across repeated trials of the same stimulus (Figure 1F-1G, Figure S1A). The pairwise relationship between odor representations in MB coding space was quantified using the correlation distance 1-*r*, where *r* is Pearson’s correlation between the vectors of KC responses to each pair of odor. Quantifying odor relationships using other metrics such as cosine distance yielded similar results (e.g., Figure S3H). When we computed the correlation distance between the KC response vector for every pairwise combination of trials in an experiment (Figure 1I, left) and reordered the distance matrix to group together trials by odor, we observed that KC responses to repeated presentations of the same odor were very strongly correlated (on-diagonal blocks, Figure 1I, right). These results demonstrate the reliability of KC odor responses across multiple presentations spanning the time course of an experiment. As expected, the pairwise correlation distance between odors reflected the qualitative similarity of their respective KC response patterns (Figure 1H-I), with visually similar activity patterns corresponding to short odor distances.

The percentage of KCs responding to each odor was similar in each MB, ranging from ~5-13% depending on the odor (Figure S1B). When compared against mean OR activity for each odor, estimated by averaging the firing rates evoked by each odor across all ORs in the Hallem dataset, KC response rate was not significantly correlated with mean ORN response strength (Figure S1B, Spearman’s rho=0.48, *p*=0.05). However, we note that the Hallem dataset underestimates ORN population responses to acids and amines since it does not include odorant receptors from the ionotropic receptor (IR) family^37^.

Overall, KCs were narrowly tuned, with most cells responding to two or fewer odors, and a significant fraction (~34%) responding to no odor in a diverse 17-odor panel (Figure S2B). However, compared against modeled KC responses (see below), observed odor responses in KCs were more broadly tuned. This result is consistent with observations that existing MB models poorly predict KC response rates to narrowly activating odors that selectively excite only one or very few ORN classes (e.g., CO_2_ or methyl salicylate, Figure S1D-E). These results confirm that KC responses are sparse and selective. They also indicate that current assumptions about MB circuit properties do not fully account for observed KC response rates to all odors, particularly for narrowly activating odors.

### Representations of odor in the MB are structured and invariant across individuals

In most circuits of the fly brain, neuronal connectivity is invariant across individuals, but the MB is distinct in that the wiring of PN inputs is probabilistic: each of the ~2000 KCs, the principal neurons of the MB, integrates input from a quasi-random subset of PNs comprising ~10% of the ~50 olfactory glomeruli in the system. As the number of possible glomerular combinations far exceeds the number of KCs in any given MB, stereotyped KC connectivity, defined by a specific set of synaptic inputs, does not exist across individual MBs. Indeed, a small set of genetically defined KCs labeled by a sparse Gal4 driver did not exhibit stereotyped odor tuning^38^. However, feedforward network models that assume random divergent connectivity between second- and third-order olfactory layers predict that, while third-order olfactory responses will be comparatively decorrelated, they will otherwise maintain relative pairwise odor relationships present in the prior layer. Thus, these models predict that the geometry of third-order olfactory representations will be invariant across different instantiations of the network (i.e., MBs), with preserved odor relationships that are predictable from the stereotyped tuning of OR inputs.

To evaluate these ideas, we compared the correlational structure of odor representations encoded in KC activity patterns in multiple flies. We found that the relationships between odors in MB representational space were indeed structured and highly invariant across individual brains (Figure 2A, S2A), consistent with prior work^3^. We evaluated the similarity of the correlational structure across MBs by comparing the rank order of odor-odor correlation distances in each MB (Figure 2B-C). The observed distribution of MB-MB correlations (Spearman’s *rho*=0.76 +/-0.06) was significantly different from shuffled controls (Figure 2D, *p* <10^-4^). Also, the correlation distance between specific pairs of odors was consistent across individual MBs: odors that evoked similar KC response patterns in one fly tended to also have similar KC responses in other flies, and the same was observed for odors that evoked dissimilar representations (Figure 2E, S2B; *p*=10^-186^, one-way ANOVA). Finally, we combined all KCs that respond to one or more odors from four different flies and clustered the KCs based on each cell’s odor response profile. This analysis identified reliable response types with specific odor tuning profiles, and each response type was found in each of the four flies (Figure 2F). For instance, KCs that are strongly co-tuned to 2-heptanone, isoamyl acetate, and pentyl acetate were reliably observed in every MB (Figure 2G). These results demonstrate that the structure of the representational space of odors is highly conserved across individual MBs in the fly.

**Figure 2:**
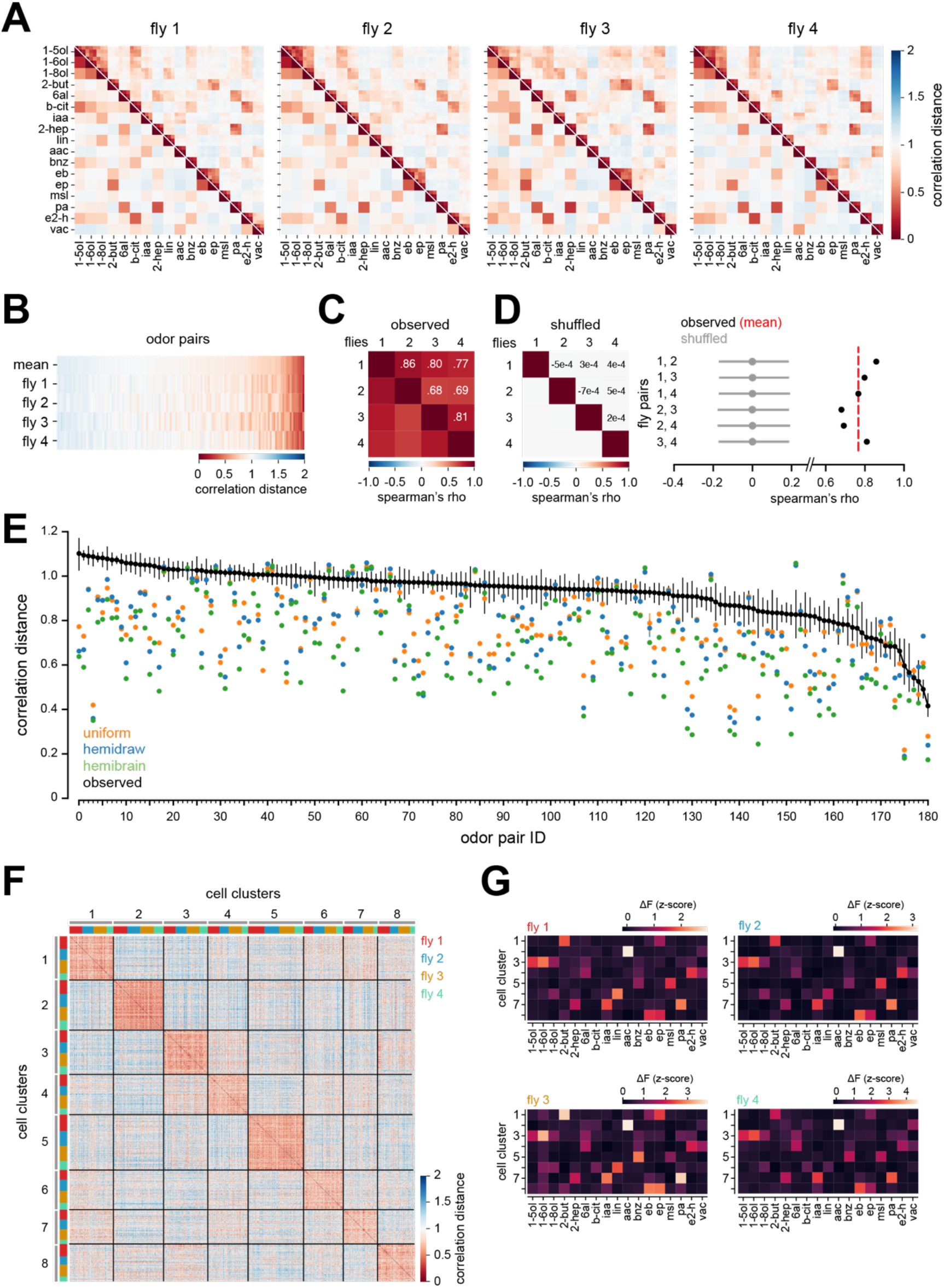
MB representational space is structured and invariant across individuals. **A)** Correlation distance matrices for four different flies showing pairwise relationships between KC population responses in individual odor trials (upper triangles) or in trial-averaged responses for each odor (lower triangle). **B)** Correlation distance between trial-averaged KC responses for every odor pair for the four flies in **A**. Odor pairs are in the same order in each row, arranged by the rank of their mean correlation distance across the four flies. Odors that evoke similar KC response patterns in one fly tend to also elicit similar response patterns in other flies. **C)** Spearman’s rank correlation between the rows of **B** (i.e., between flies), evaluating the similarity of the rank ordering of odor pairs between flies according to their representational distance in KCs. **D)** Left: same as **C**, but for shuffled data in which the odor labels were randomly permuted for the responses of individual flies. The matrix shows the mean Spearman’s correlation across 10,000 shuffles. Right: observed Spearman’s correlation (black) and the mean and 95% CI of the Spearman’s correlation across 10,000 shuffles (grey) for each fly pair. Red dotted line marks the mean observed Spearman’s correlation (rho=0.76). **E)** Correlation distance (mean and 95% CI) between KC responses for each odor pair, averaged across all flies in which the odor pair was sampled (*n*=3-22, see Supplemental Table 1). Each unique odor pair was assigned a reference ID (see Supplemental Table 1). A one-way ANOVA showed there was a significant difference in the odor-odor correlation distance between different odor pairs (F statistic=12.6, *p*=10^-180^), consistent with odor-odor relationships being reliable across MBs in different flies. The correlation distance (mean and 95% CI across 100 model MBs) between predicted KC responses for odor pairs in the uniform, hemidraw, and hemibrain models are plotted for reference (see Figure 3). **F)** Matrix of pairwise correlation distances between odor response profiles of every KC in the four flies in **A** that responded to at least one odor. The distance matrix was ordered by spectral clustering on the mean odor response vector of each cell. Each response cluster contained KCs from every fly. **G)** Mean KC tuning profiles of each cluster, computed across KCs in each cluster in each fly. KCs with conserved odor tuning profiles are found in every MB.

### Odor distances in MB coding space diverge from odor relationships predicted by a random feedforward network model

To compare the observed structure of MB representational space to what is predicted by a feedforward random network, we modeled KC population responses by adapting a previously described, biologically plausible spiking model of the fly olfactory circuit^30^ (Figure 3A). PN responses, modeled from OR firing rates in the Hallem dataset^32^, were used as input to a population of 2000 spiking KCs. KCs were modeled as leaky integrate-and-fire units with a small number of input sites, typically ~5-7 dendritic claws, that each receive input from a single PN bouton. Under assumptions of random feedforward connectivity, the matrix of PN-KC connections in each model instantiation was created by assigning each KC claw a single glomerular input, with all glomeruli having an equal likelihood of being drawn (“uniform” model). The inputs to the model were the firing rates of approximately half of fly ORs to 109 odors from the Hallem dataset. KC activity was normalized by global feedback inhibition from the GABAergic APL. KC spiking thresholds and APL-KC inhibitory weights were tuned to achieve a mean KC response rate of ~10% across odors (see Methods), which matches experimental observations.

**Figure 3:**
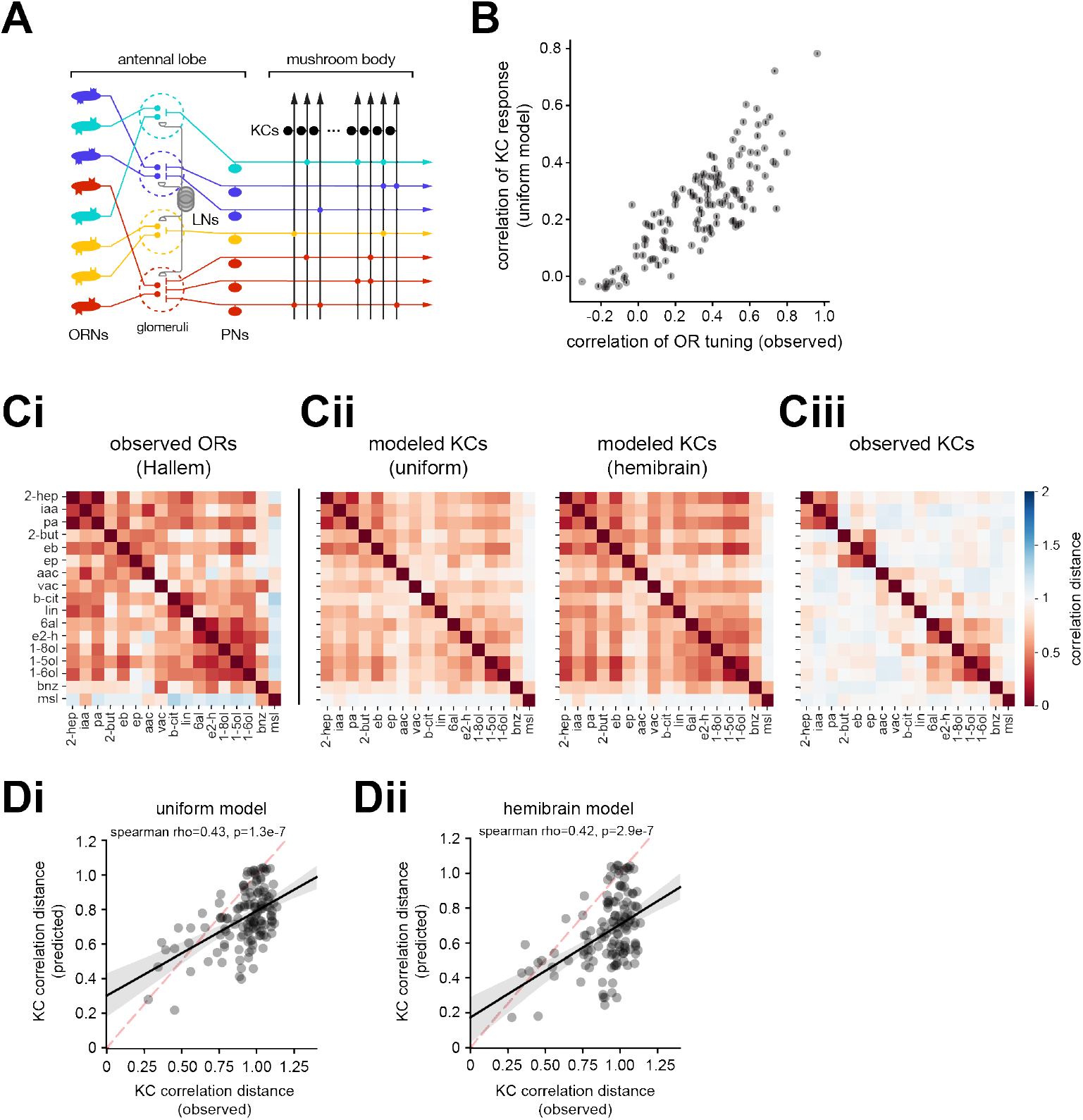
Odor relationships in MB representational space deviate from predicted relationships in a random feedforward network. **A)** Schematic of the fly olfactory network. Not depicted is a single GABAergic neuron (APL) in the MB that mediates feedback inhibition among KCs. **B)** Comparison of relationships between predicted KC representations (uniform model) and observed OR tuning profiles for each odor pair (grey symbols). Error bars are 95% CI of KC correlation for each odor pair across 100 model MBs; each simulated MB has an independently generated PN-KC connectivity matrix drawn under assumptions of uniform input (each glomerulus drawn with equal probability). **C)** Matrix of pairwise correlation distances between OR tuning profiles from the Hallem dataset **(Ci)**; predicted KC responses in the uniform model (**Cii**, left) or hemibrain model (**Cii**, right); or observed KC responses (**Ciii**) for 17 odors. **D)** Comparison of correlation distances for each odor pair between observed KC responses for each odor pair and predicted KC responses in the uniform (**Di**) and hemibrain (**Dii**) models. Each symbol is an odor pair.

The correlation between modeled KC responses for specific odor pairs was indeed consistent across 100 simulated MBs with different, independently drawn random PN-KC connectivity matrices (Figure 3B, error bars are 95% CI across simulations). However, the observed structure of odor representations in KC coding space was only partially predicted by the feedforward random model, with the observed relationship between odors deviating significantly from model predictions for many odor pairs (Figure 2E, 3C, Di, S3H). In particular, observed pairwise odor relationships were overall more decorrelated than those predicted from modeled KC populations (Figure 3Di). Adjusting fit parameters to yield a lower mean KC response rate of 5% across odors did not appreciably affect the systematic overprediction of the degree of correlation between odor representations, nor did it affect the rank order of predicted odor-odor correlation distances (data not shown).

Recent analysis of the global structure of glomerular input sampling by third-order olfactory neurons (Figure S3A) using large-scale EM-level reconstructions of part of the *Drosophila* brain at synaptic resolution revealed that the wiring of PN inputs to KCs is not fully random^18,19^, as was previously believed. Indeed, analysis of the complete matrix of PN-KC connectivity from two independently reconstructed fly brains confirmed that KCs sample particular combinations of PN inputs at a higher rate than is expected based on their numerical frequency, and the structure of this biased input onto KCs is similar between two MBs from different flies^19^ (Figure S3B-E). We also observed that the glomerular input structure to KCs bears significant similarity to the structure of glomerular input to third-order olfactory neurons in the lateral horn (Figure S3E), a brain region dedicated to innate odor processing in which neurons have stereotyped connectivity and tuning^39,40^.

To assess the possible impact of structured input on MB odor representations, we predicted KC odor responses in the model under conditions in which the PN-KC connectivity matrix was drawn according to the observed frequency of PN boutons corresponding to each glomerulus in the hemibrain dataset (“hemidraw”) or in which the experimentally reconstructed hemibrain PN-KC connectivity matrix was directly implemented (“hemibrain”) (see Methods). Neither adjustment to PN-KC connectivity improved predictions of observed KC odor relationships (Figure 3C, 3Dii, S3F-H); in fact, predicted responses for most odor pairs tended towards being more correlated compared to the uniform model. These results indicate that observed biases in PN-KC connectivity are unlikely to account for the differences between observed and predicted KC odor responses; they suggest additional sources of structure are present in the olfactory circuit that mediate the observed transformation of odor representations from the periphery to the MB.

### Reorganization of representations of odor in the MB around odor relationships in natural sources

Comparisons of odor representations in the OR input layer and in the MB showed that, although the representations of most pairs of odors were substantially decorrelated, as expected, some odor pairs were comparatively less decorrelated between the OR and MB layers. This transformation resulted in a significant regrouping of odor relationships in OR versus MB coding space. For instance, the odors 2-heptanone, isoamyl acetate, and pentyl acetate emerged as a cluster with similar KC representations, distinct from other odors in the panel, whereas the representation of each of these odors is similar to many others at the level of its OR representation (Figure 3Ci, iii).

To better understand the functional implications of this reorganization, we asked how odor representations at different stages of olfactory processing relate to the properties of the odors, focusing in particular on their chemical properties and on how they are correlated across natural odor sources. For each odor, we computed molecular and physicochemical descriptors using Mordred, an open-source molecular descriptor calculation software^41^. Since many descriptors are highly correlated across odors, we identified a reduced set of 570 molecular descriptors that captured the chemical relationships between odors equivalently to the full set of descriptors (Supplemental Table S3).

*D. melanogaster* is an ecological generalist and human commensal^42^; as a starting point for understanding the structure of natural odor space for *Drosophila,* we used a database of the headspace volatile profiles for many food odor sources, compiled from 4407 published chemical datasets primarily from the food and flavor science literature (Figure 4A). The odor sources are biased towards fruits, plants, and vegetables, but also includes other common human foods such as alcoholic beverages, meat, and dairy products. The database comprises thousands of samples measured from 887 types of natural sources (apple, tomato, wine, etc.) and contains over 8,000 monomolecular volatiles. Like other datasets profiling the chemical volatiles emitted from natural sources^43^, the majority of odorants occur sparsely in a small number of sources, though a significant minority of odors are present broadly across many sources (Figure 4A).

**Figure 4:**
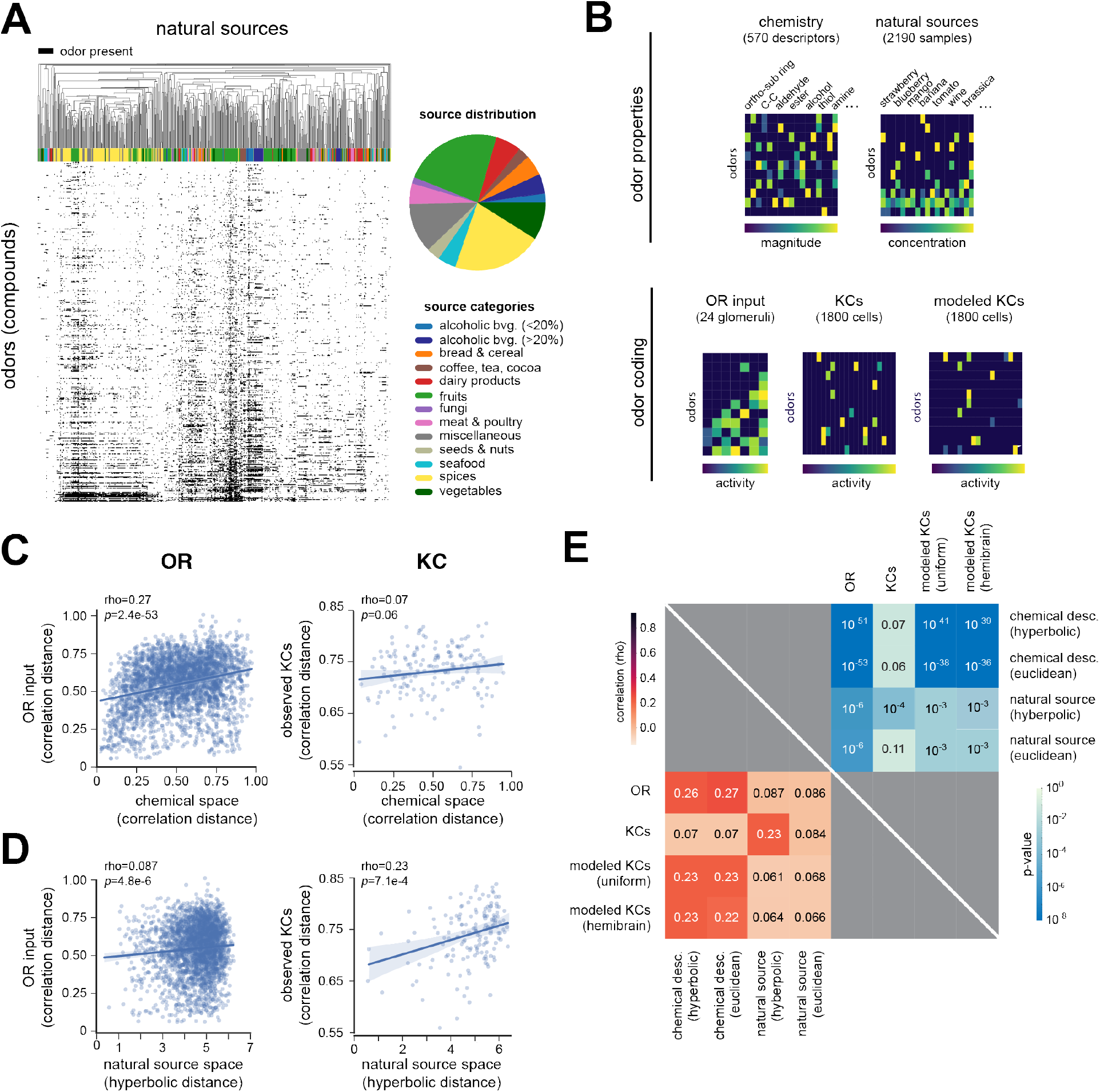
Reorganization of odor representations in the fly MB reflects odor relationships in natural sources. **A)** Distribution of odors (monomolecular compounds, rows) in a literature-based database of the volatile headspace composition of many natural food odor sources (columns). The pie chart shows the distribution of odor sources in each food category. **B)** Schematic summary of datasets for comparing the properties of odors (top) with their different neural representations (bottom). **C)** Comparisons of odor-odor correlation distances in OR (left) or KC (right) coding space with their distances in chemical descriptor space. **D)** Comparisons of odor-odor correlation distances in OR (left) or KC (right) coding space with their embedded distances in a hyperbolic model of natural source space. **E)** Summary of correlation (lower triangle) and *p*-values (upper triangle) for comparisons between observed OR, observed KC, or modeled KC (uniform or hemibrain) odor relationships and odor-odor relationships in chemical or natural source space.

Recently, we showed that the use of hyperbolic coordinates to embed the concentrations of individual monomolecular odorants as they occur in natural odor sources better captures odor relationships in the space of natural sources, compared to embeddings that use Euclidean metrics^12,45^. Thus, to describe the similarity in distribution across natural odor sources for a given pair of odors, we started with a normalized correlation coefficient computed between odor abundances across different natural sources. Based on these distances, we performed nonlinear dimensionality reduction using different curved and flat metrics^64^. The method automatically adjusts the curvature of the embedding space and selects the best fitting dimension based on the Bayesian Information Criterion^44^. We found that the natural odor abundance data was best described by a three-dimensional hyperbolic space with a negative curvature of −5.12. This observation mirrors previous results using a more limited dataset of natural fruit or flower odor sources in which hyperbolic geometry also provided a significantly better fit of the natural source space than the standard Euclidean geometry^12,45^. The intuitive explanation for these results is that hyperbolic spaces provide good descriptions of the structure of natural odor spaces because they arise as continuous approximations to tree-like hierarchical networks. In the case of natural source data, the hierarchical relationships are hypothesized to reflect dependencies produced by biochemical and metabolic pathways acting in plants and other food sources, including in associated microbes like bacteria and fungi. In contrast, nonlinear dimensionality reduction of odor distances computed from chemical descriptors resulted in a hyperbolic embedding with higher dimension (dim=6) and much reduced curvature (0.05) that did not provide a better fit of the dataset than a Euclidean model. Since odor distances in the hyperbolic embedding better captured the correlational structure of abundances of odors across natural sources compared to equivalent distances computed from a Euclidean embedding, we used hyperbolic distances as a metric of odor relationships in natural source space.

For each pair of odors in the dataset, we compared their representational distance in OR or KC coding space with their relationship in chemical descriptor space or natural source space (Figure 4B). We found that pairwise odor distances in OR coding space were better correlated with their distances in chemical descriptor space (*rho*=0.27, *p*=2.4e-53) as compared with their distances in natural source space (*rho* =0.087, *p* =4.8e-6) (Figure 4C-D). The converse was true for odor distances in KC coding space: pairwise distances in KC coding space were better correlated with their distances in natural source space (*rho* =0.23, *p* =7.1e-4) compared to their distances in chemical descriptor space (*rho* =0.07, p=0.06) (Figure 4C-D). The correlation to distances in KC coding space was observed only for natural source odor relationships quantified by the hyperbolic embedding of natural source space, but not by Euclidean distance (Figure 4E), suggesting that capturing hierarchical relationships in the dataset is important and that the low-dimensional embedding helps to de-noise the data. We also evaluated the relationship of odor properties to odor distances in the modeled KC coding space under assumptions of unstructured PN-KC connectivity. In contrast to the observed KC distances between odors, modeled KC distances were better correlated with odor distances in chemical descriptor space (*rho*=0.23, *p*=1.4e-38) compared to hyperbolic odor distances in natural source space (*rho*=0.061, *p*=8.4e-3) (Figure 4E). This result is expected since, under assumptions of unstructured connectivity in the olfactory circuit, the relationship of odors in KC coding space stems from their relationships in OR input space. Similar results were observed for odor relationships computed from KC responses predicted using the hemibrain circuit model, indicating that the observed degree of glomerular input bias to KCs in the hemibrain does not explain the reformatting of odor relationships in the MB. Overall, these data indicate that the realignment of odor representation with natural source relationships that we observed in KCs cannot be derived simply by random resampling of OR responses.

To determine if the relationships between odor representations and odor properties were driven by only a small number of odor pairs in the dataset, we recomputed the correlations between OR or KC representational distance and chemical descriptor or natural source distance using 100 subsamples comprising 75% of the odor pairs, dropping out a random 25% of the odor pairs in each resampling. This analysis confirmed that odor distances in OR space were more correlated with distances in chemical descriptor space as compared to in natural source space, and vice versa for odor distances in KC space. These findings are consistent with a reorganization of the fly olfactory code between the periphery and the MB from encoding the chemical or structural relationships between odors to reflecting the odor relationships in complex mixtures arising from natural sources.

### Odor relationships are restructured across successive stages of olfactory processing

To better understand how representations of odor are reformatted from the periphery to the MB, we used functional imaging to measure odor-evoked patterns of activity at each successive stage of processing in the olfactory circuit. While delivering odors to the antennae of the fly, we volumetrically imaged from all ORN axon terminals (labeled by the *pebbled-Gal4* driver) or from ~70% of PN dendrites (labeled by the *GH146-Gal4* driver) in the antennal lobe^46^, where the neurites from these cell populations are stereotypically organized into glomerular compartments with characteristic size and position (Figure 5A-B). Population imaging from the terminals of all ORN classes allowed odor relationships to be assessed in the complete OR input space to the olfactory system, whereas the Hallem dataset measures the tuning of only approximately half of all fly ORs (Supplemental Figure S4B). ROIs corresponding to individual glomeruli were manually segmented from movies of odor responses in ORN axon or PN dendrites and their odor response profiles extracted. To evaluate patterns of odor-evoked PN output, we volumetrically imaged from the axonal boutons of ~70% of PNs (labeled by the *GH146-Gal4* driver) in the calyx of the MB, where they synapse onto the sites of KC input (claws) (Figure 5A-B). ROIs corresponding to PN boutons were automatically segmented using Suite2P; visual inspection confirmed that > 95% of ROIs reliably corresponded to single PN boutons. Odor relationships were invariant across the brains of different individuals at each of these earlier stages of olfactory processing (Figure S4A), consistent with the stereotyped connectivity of the antennal lobe.

**Figure 5:**
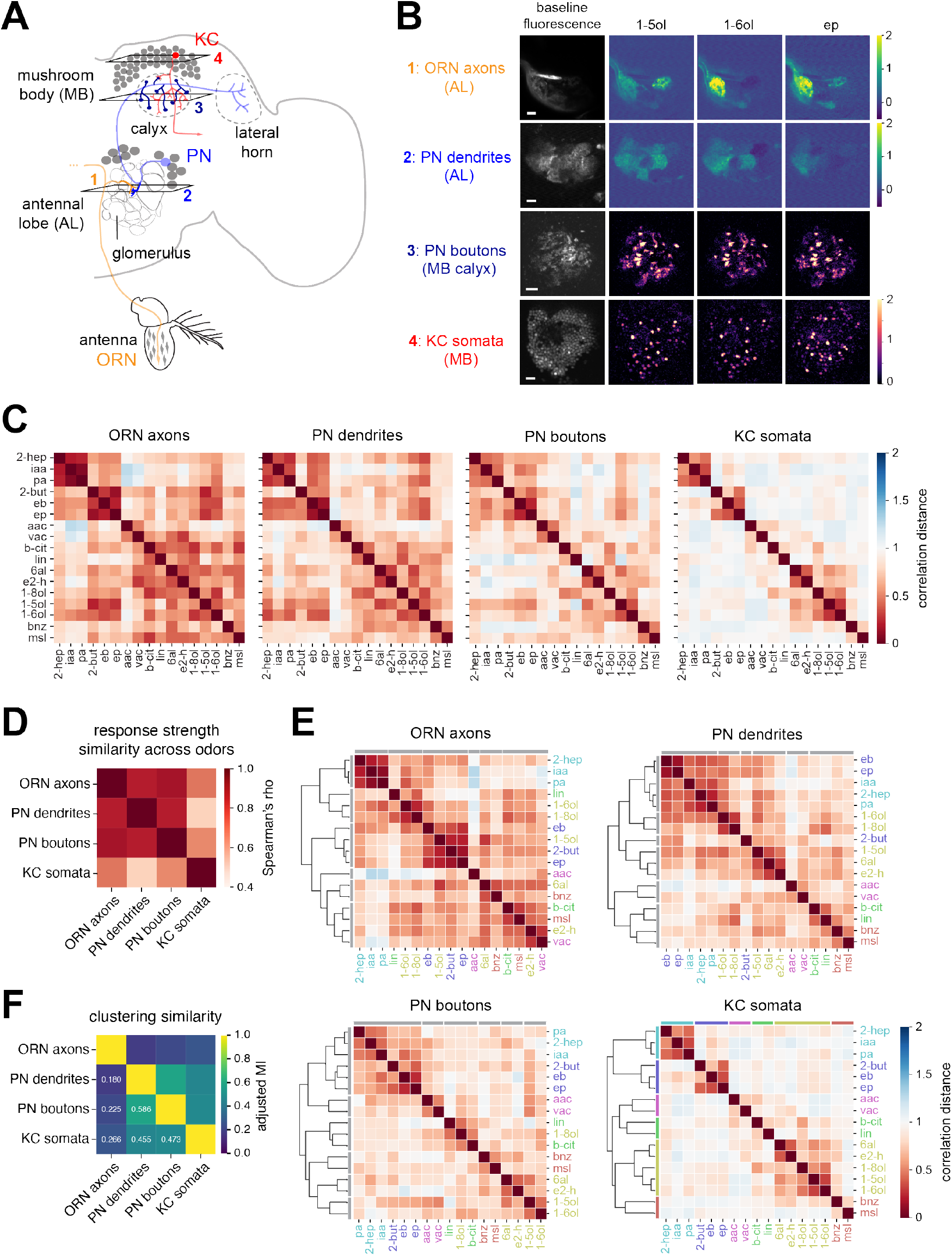
Reformatting of odor representations across four stages of olfactory processing. **A)** Anatomical schematic of fly olfactory circuit. ORN axon terminals are organized in a stereotyped glomerular map in the AL. Each uniglomerular PN has dendrites in a single AL glomerulus and projects to the calyx of the MB, where its axonal arbor terminates in multiple boutons that each synapse with KC input claws. **B)** Representative baseline fluorescence and peak △F/F heatmaps of odor-evoked calcium signals in ORN axon terminals, PN dendrites, PN boutons, and KC somata to 1-pentanol, 1-hexanol, and ethyl propionate. KC data are reproduced from Figure 1. Scale bar, 10 μm. Genotypes: ORN axons are imaged in *pebbled-Gal4, UAS-GCaMP6f*; +; *GH146-QF, QUAS-tdTomato*. PN dendrites and PN boutons are imaged in *GH146-Gal4/20xUAS-IVS-jGCaMP8m* (II). **C)** Matrix of mean pairwise correlation distances (averaged across flies) of trial-averaged odor representations in ORN axons (*n* = 4 flies), PN dendrites (*n* = 6 flies), PN boutons (*n* = 6 flies), and KC somata (*n* = 4 flies) to a panel of 17 odors. Odors are displayed in the same order in each matrix according to the clustering of their representations in KC representational space. **D)** Spearman’s correlations measuring the similarity of rank ordering of odor response strengths at each stage of processing. **E)** Hierarchical clustering of distance matrices in **C** showing best odor groupings in each representational space. **F)** Pairwise adjusted mutual information (MI) score evaluating the similarity of clustering of odor representations in each representational space. The adjusted MI has a value of 1 when two clusterings are identical, and a value of 0 when the MI between two clustering is the value expected due to chance.

Comparing the pairwise correlation distances between odor representations in each layer of the olfactory circuit, odor representations were progressively decorrelated at each feedforward stage of processing (Figure 5C), with the mean and median odor-odor correlation becoming systematically lower in each successive layer (Figure S4D). The largest decorrelation occurred between PN boutons and KC somata, consistent with the divergent expansion of the circuit at this synaptic step. However, the rank order of odor-odor distances was not maintained across layers (ORN terminals versus KC somata, Spearman’s rho=0.48), indicating that odor relationships are transformed beyond a simple linear scaling between layers. For comparison, in a model that assumes unstructured PN input to KCs and uniform inhibition in the AL and MB (“uniform”), the Spearman’s correlation between odor relationships in the input OR layer and in modeled KCs is rho=0.83. Furthermore, whereas the relative levels of odor-evoked activity in PNs dendrites or boutons was well-predicted by levels of overall ORN input to the circuit, odor-evoked KC response rates (and mean activity) were comparatively less well-predicted (Figure 5D, S4Ci-iv, and data not shown). These observations all point towards a non-linear reformatting of odors in the MB representational space that alters the rank ordering of odor relationships compared to earlier stages of coding.

We examined odor relationships at each step of processing using hierarchical clustering of odor representations in each neural population, in order to better understand the origins of odor relationships in MB coding space. For instance, one of the strongest odor clutsers at the level of input representations – 2-heptanone, isoamyl acetate, and pentyl acetate – was maintained in each subsequent stage of processing through to the KCs (Figure 5C, E). These odors share a high degree of chemical similarity as well as co-occurrence in natural sources, which may explain their robust grouping. However, other odor representations that were similarly well-correlated and clustered in OR space (e.g., 1-pentanol and 2-butanone) were selectively decorrelated in KC coding space, with each odor becoming reassigned to distinct, non-overlapping groups of odors in KC coding space (Figure 5C, E). Additionally, the representations for some odor pairs were observed to become reliably *more* similar at later stages of olfactory processing. For instance, the pairwise correlation distances between 1-pentanol and 1-hexanol was actually shorter in KC coding space as compared to in ORN axon, PN dendrite, or PN bouton coding space.

Typically, these transformations in the representational geometry among odors occur progressively at each successive stage of processing, with the largest change occurring between PN boutons and KC somata (Figure 5C). However, we observed significant reorganization of odor representations even between the inputs and outputs of the same cell population, PNs; odor relationships encoded in PNs boutons better reflect odor relationships in KC coding space compared to PN dendrites (Figure 5C, E). The nonlinear reformatting of odor relationships across successive stages of olfactory coding indicates that prevailing models of olfactory networks that assume global or uniform processing across neurons are incomplete, and additional sources of structure exist in the fly olfactory circuit that mediate non-uniform interactions between olfactory coding units (e.g., neurons or glomeruli).

## DISCUSSION

We show that representations of odor are structured and invariant across MBs in individual flies, and that the structure of MB odor coding space is only partially predicted by models that assume random sampling of olfactory glomerular inputs by KCs. The latter is true even after adjusting the MB model to account for the over- or under-representation of PN boutons from specific glomeruli (“hemidraw” model) or the amount of over- and under-convergence of PN inputs from specific sets of glomeruli onto KCs (“hemibrain” model). Thus, certain assumptions of olfactory system architecture – for instance, of uniform strengths of unitary feedforward synapses, uniform spiking thresholds, or nonselective, global inhibition in the AL or MB – are likely oversimplifications, motivating a search for source(s) of structured interactions between glomeruli or neurons in the olfactory circuit.

We demonstrate a significant transformation of the fly olfactory code between the periphery and the MB, in which the encoding of odors by ORs better reflects the chemical relationships between odors and the encoding of odors in the MB better reflects the distributions of odors across behaviorally relevant natural sources. That representations at the olfactory periphery better capture odor relationships in terms of their chemical or molecular properties is perhaps unsurprising, since odor-OR interactions are governed by the structural features of the odor. The reorganization of MB odor representations to better correlate with the relationships of these odors in natural source space may reflect the progressive transformation of odor representations in the olfactory network to encode latent variables that relate more directly to behavioral value or perception.

Odor distances in the hyperbolic embedding of natural odor mixtures reflect correlations between volatiles in the headspace profiles across natural sources that arise from conserved metabolic pathways^45^. The transformation of odor representations in the MB is predicted to facilitate the perceptual compression of odors that have shorter metabolic tree distances in the hierarchical organization of natural odor mixtures relative to their chemical similarity. Consistent with this idea, a recent preprint reports that distances between odors in a neural network embedding trained on human olfactory perceptual labels are correlated with the metabolic distance between odors in experimentally elucidated biochemical pathways^47^.

The specific structure of odor relationships was largely invariant across individual brains at each stage of olfactory processing (Figure 2), even in the MB where PN boutons connect probabilistically with KCs. An important open question is the extent to which the invariance of MB representational space may arise from genetically specified developmental processes (sculpted by evolution) or from activity-dependent processes that reflect shared olfactory experience^2^. Unlike their mammalian analogues in piriform cortex, KCs are not strongly connected through recurrent excitation, although the degree to which PN-KC synapses or APL-KC synapses may be regulated by experience is not well understood. We note that the structure of PN overconvergence onto KCs bears significant similarity to the structure of PN input to third-order olfactory neurons in the lateral horn (LH) (Figure S3E). LH neurons are stereotyped in their anatomical connectivity and odor tuning and are believed to mediate innate olfactory behaviors. The similarity in PN input structure between the MB and LH points to the likely ethological significance of these over-represented glomerular combinations. It also raises the possibility that the LH may have additional roles in shaping odor representations in the MB. For instance, neurons downstream of the LH send centrifugal input to the MB calyx^48^ which could contribute to the remapping of odor representations in PN boutons. Other possible sources of structure in the circuit are selective inhibition from the APL or MB-C1 neurons in the calyx, or non-uniform PN-KC synaptic weights. Understanding the specific circuit mechanisms that shape the reorganization of odor representations in the MB will be important for understanding how odor relationships in natural sources become reflected in the olfactory code.

Theoretical studies of divergent expansive cerebellum-like circuits such as the MB emphasize the computational benefits of random networks for maximizing coding capacity and promoting the separability and discriminability of representations. However, structured networks can correlate odor representations and promote generalization of odors along important directions of odor space, for instance, related to the relationships of odorant molecules in behaviorally salient natural odor sources. Our results suggest a revision of classic formulations of cerebellar-like network architectures is warranted at least in some systems: the MB may trade-off the capacity to maximally decorrelate activity patterns in parts of representational space for an increase in coding capacity and robustness in a part of representational space of particular ethological significance^49^. The recoding of odors in the MB to reflect relationships in natural sources would predict greater generalization to odors with similar distributions in the environment, facilitating the decoding of natural source identity in noisy or ambiguous situations.

## METHODS

### Experimental model

*Drosophila melanogaster* were raised on a 12:12 light:dark cycle at 25°C and 70% relative humidity on cornmeal/molasses food containing: water (17.8 l), agar (136 g), cornmeal (1335.4 g), yeast (540 g), sucrose (320 g), molasses (1.64 l), CaCl2 (12.5 g), sodium tartrate (150 g), tegosept (18.45 g), 95% ethanol (153.3 ml) and propionic acid (91.5 ml). All experiments were performed in female flies aged 3-10 days post-eclosion. Unless otherwise noted, the transgenes in this study were acquired from the Bloomington Drosophila Stock Center (BDSC) and have been previously characterized as follows: *pebbled-Gal4* (X) directs expression in all ORNs^50^ (RRID:BDSC_80570); *GH146-Gal4* (II) directs expression in ~70% of PNs^51^ (RRID:BDSC_30026); *GH146-QF, QUAS-tdTomato* (III) expresses the red-fluorescent protein tdTomato in ~70% of PNs^52^ (RRID:BDSC_30037); *UAS-IVS-jGCaMP8m* (II) expresses the calcium indicator jGCaMP8m^53^ in a Gal4-dependent manner (RRID:BDSC_92591); *OK107-Gal4* (IV) directs expression in all KCs^33^ (RRID:BDSC_854); *UAS-OpGCaMP6f* (X) was from B. D. Pfeiffer and D. J. Anderson (Caltech, Pasadena, CA) and expresses the calcium indicator codon-optimized GCaMP6f^54^ in a Gal4-dependent manner; and *UAS-nls-OpGCaMP6s-p10 (III)* was from H. Chiu and D. J. Anderson (Caltech, Pasadena, CA) and expresses nuclear-localized, codon-optimized OpGCaMP6s^35^ in a Gal4-dependent manner.

### Odor stimuli

Odors were delivered essentially as previously described^55^. A custom-built multi-channel olfactometer delivered a constant 2 L/min stream of charcoal-filtered air. A 3-way solenoid valve directed 200 mL/min of this flow either through a 20-ml glass vial containing 2-ml of odor solution (valve open) or an equivalent vial containing 2-ml of the solvent. Air flow was controlled using mass flow controllers (MC series, Alicat Scientific, Tucson, AZ). The 200 ml/min control or odor streams were carried by tubing of matched lengths and rejoined the carrier stream at the same point along the carrier tube, approximately 10 cm from the fly. The terminal end of the carrier tube had an inner diameter ~8mm and was ~1 cm away from the fly.

Odor concentrations refer to the v/v dilution factor of the odor solution in the vial. The concentration of odor in the headspace is further diluted 10-fold in air prior to reaching the fly. Unless otherwise indicated, all odors in this study were presented from a 10^-3^ (0.1%) dilution in paraffin oil (J.T. Baker, VWR #JTS894), with the exception of movies collected from one fly, in which odors were diluted to 10^-2^ (1%).

### Two-photon calcium imaging

Volumetric, in vivo functional calcium imaging was performed essentially as previously described^46,55^. After a brief period of cold anesthesia (<20 s), the fly was head-fixed, and the cuticle, fat, and air sacs were removed to expose the brain region of interest. Two-photon GCaMP6f or GCaMP8m fluorescence was excited with 925 nm light from a Mai Tai DeepSee laser (Spectra-Physics, Santa Clara, CA). Images were acquired with an Olympus 20X/1.0 numerical aperture objective (XLUMPLFLN20XW) driven by a piezo motor that enabled fast z-scanning, and the collection filter was centered at 525 nm with a 50 nm bandwidth. Experiments were conducted at room temperature (~22°C). The brain was constantly perfused by gravity flow with saline containing (in mM): 103 NaCl, 3 KCl, 5 N-Tris(hydroxymethyl)methyl-2-aminoethane-sulfonic acid, 8 trehalose, 10 glucose, 26 NaHCO3, 1 NaH2PO4, 1.5 CaCl2, and 4 MgCl2 (pH 7.3, osmolarity adjusted to 270– 275 mOsm). The saline was bubbled with 95% O_2_/5% C_O_2 and circulated in the bath at ~2-3 ml min^-1^.

#### AL imaging

Flies were head-fixed with the dorsal surface of the head approximately parallel to the imaging plane; the dorsal cuticle was removed, and the antennal lobes were exposed. The antennae were snugly secured below the imaging chamber, keeping them dry and responsive to odors. ORN axon terminals and PN dendrites were imaged with a galvo-galvo scanning system (Thorlabs Bergamo). Flies were imaged from the dorsal side, with 5 planes spaced 12 μm apart. The depth of the first imaging plane was chosen to maximize the number of glomeruli visible across the 5 planes. Images were acquired at a resolution of 192×192 pixels, with typical fields-of-view of ~90-100μm^2^ and volumetric sampling rates of ~1 Hz through the antennal lobe. For ORN axon and PN dendrite experiments, odors were presented in a pseudorandom order, with the three trials of each odor being presented contiguously. Trials consisted of a 7 s baseline recording, the odor pulse (2s for ORN experiments and 3s for PN experiments), and a 20 s post-stimulus period.

#### MB imaging

Flies were head fixed with the head tilted acutely downward, rotated ~70° from its normal resting position. The posterior plate of the head was approximately parallel to the imaging plane, and the antennae were dry underneath the imaging platform. The entire perimeter of the head capsule was stabilized to the imaging platform with glue, and the proboscis and legs were immobilized to minimize motion. KC somata and PN boutons were imaged with a galvo/resonance scanning system (Thorlabs Bergamo), with the exception of one fly in which KC somata were imaged on a galvo-galvo scanning system at a volumetric sampling rate of 0.55 Hz. For PN bouton experiments, odors were presented in a pseudorandom order, with the three trials of each odor being presented contiguously. Trials consisted of a 7 s baseline recording, a 3s odor pulse, and a 20 s post-stimulus period, with stimulus onset occurring every 30s. For KC somata experiments, stimulus order was pseudorandomized such that all odors were presented before any were repeated; thus, repetitions of a given odor usually did not occur on consecutive trials. If paraffin was included as an odor stimulus, it was always presented first in each repetition block. Trials consisted of a 13 s baseline recording, a 2 s odor pulse, and a 45 s post-stimulus period, with stimulus onset occurring every 60s.

For resonance imaging of KC somata, movies were collected at a frame rate of ~60 Hz at 256 x 256 pixels in fast-z mode. The field-of-view was ~150 μm x 150 μm giving a pixel size of ~0.6 μm/pixel. The bulk of the KC somata cluster was captured in 16 planes (and 4 flyback frames) collected 2 μm apart, spanning a ~30 μm depth through the MB. This resulted in a volumetric sampling rate of ~3 Hz. For a subset of experiments, KCs were sampled using fewer imaging planes (eleven) with a z-step size of 3 μm. Since odor response rates and odor-odor relationships were similar with these imaging parameters, KC somata datasets were combined. For imaging of PN boutons in the MB calyx, movies were collected at a frame rate of ~60 Hz at a resolution of 256 x 256 pixels in fast-z mode. The field of view was ~75 μm x ~75 μm yielding a pixel size of ~0.3 μm/pixel. The full population of labeled PN boutons in the calyx was captured using 8 planes (and 2 flyback frames) spaced ~3 μm apart, spanning a ~21 μm depth in *z*, giving a volumetric sampling rate of ~6 Hz.

### Image analysis

#### AL imaging

Image analysis was performed using custom Python scripts (https://github.com/ejhonglab/al_analysis). Motion correction was performed separately in each plane using the registration module in Suite2p. ROIs corresponding to individual glomeruli were manually defined in each imaging plane in Fiji^56^, using a combination of the resting fluorescence and per-trial responses to visualize the glomerular boundaries. Responses (ΔF/F) were calculated using a 6 s baseline (F) that ended 1 s prior to nominal odor onset in each trial. The odor response was quantified as the mean ΔF/F during the 2 s after odor onset. Odor tuning of identified glomeruli, assigned using a combination of anatomical position, size, shape, and responses to a diagnostic panel of 10 narrowly activating odor stimuli, matched previous descriptions^32,57,58^.

#### MB imaging

##### Motion correction

For PN bouton and KC somata imaging, an initial round of 3D motion correction was performed with the ‘NormCorre’ algorithm, using either the Matlab implementation^59^ (https://github.com/flatironinstitute/NoRMCorre) or Python implementation from the calcium imaging analysis library ‘CaImAn’^60^ (https://github.com/flatironinstitute/CaImAn). As necessary, planes were dropped following motion correction due to field-of-view drift; typically the most superficial or deepest planes were dropped if they were not consistently captured through the entire recording.

##### Source extraction

For movies of PN boutons and KC somata collected in the MB, source extraction was carried out using either Suite2p^36^ or CaImAn^60^. For all PN bouton, and for KC somata data collected at a z-step of 2 μm, Suite2p was used to perform source extraction on a plane-by-plane basis. ROIs corresponding to PN boutons and odor-evoked signals were extracted in Suite2p’s *functional* mode, using the activity-based algorithm. KC ROIs and signals were extracted in Suite2p’s *anatomical* mode, using the ‘cellpose’ model to perform anatomical segmentation on the mean (time-averaged) image. Extracted fluorescence traces were neuropil-corrected (F_corrected_ = F – 0.7 *F_neuropil_) and normalized by z-scoring over time. The ‘rastermap’ function^61^ was used to visualize F_corrected_ for all extracted components, with cells sorted to cluster those with similar patterns of activity. Stimulus-responsive clusters were manually selected. For PN bouton movies, this step was used to filter out spurious ROIs – only components belonging to stimulus-responsive clusters were included in subsequent analyses. For KC movies, the relatively high level of baseline fluorescence in combination with the use of a nuclear-targeted calcium indicator resulted in very high-quality cellular segmentation, with very few (if any) spurious ROIs. Figure 1G shows the responses of all extracted cells from one representative experiment, with cells (rows) corresponding to stimulus-responsive clusters displayed at the top of the matrix, and non-responder cell clusters shown at the bottom.

CaImAn-MATLAB was used to perform 3D source extraction on KC somata data collected at a z-step size of 3 μm. For each KC, a single ROI was extracted, which was roughly spheroid and spanned multiple planes. The fluorescence baseline F0 was computed at each timepoint by taking a fixed percentile (ranging from 20-40%) of a rolling 30 s time window. Extracted raw calcium signals were detrended and normalized to baseline fluorescence F0 to compute ΔF/F responses. All components extracted by CaImAn were included in subsequent analyses. Between ~1500-1900 KCs and ~500-800 PN bouton ROIs were extracted per movie.

##### Quantification of odor response

For Suite2p extracted signals, ROI response strength in a single trial was computed by averaging the signal over an expected response peak time window (0.25-2s post-stimulus onset for PN boutons, 2-8s post-stimulus for KCs), and subtracting the mean pre-stimulus baseline (initiating 5s pre-stimulus for PN boutons, and 10s pre-stimulus onset for KC soma). For CalmAn-extracted KC signals, the response strength for each trial was calculated by subtracting the median 10s pre-stimulus baseline from the mean 15s post-stimulus ΔF/F signal. For all datasets, mean odor response strength was computed by averaging across all trials in which the odor was presented.

#### KC response rate

KC response rates (Supplementary Figure S1) were computed from datasets in which source extraction was performed using Suite2p, applying the ‘cellpose’ model to perform anatomical segmentation. All extracted components were used in the analysis of response strength and response breadth, including ‘silent’ cells that did not respond to any stimulus in the odor panel. A cell was considered a ‘responder’ if its trial-averaged response to a given odor exceeded a fixed threshold. The response threshold was determined separately for each dataset using ethyl propionate (a consistently strongly activating odor) as a reference, in order to adjust for small differences in responsiveness of different experimental preparations. The KC response rate to ethyl propionate was computed over a range of thresholds, with a step size of 0.05 between threshold values. Threshold values resulting in a 12-15% ethyl propionate response rate were selected. The median of these values was chosen as the final response threshold and was used to compute response rates for the other odors in the dataset.

#### Analysis of the representational space of odors

##### Distances between population representations of odor

The correlation distance between the population representation of two odors was computed as 1-*r*, where *r* is Pearson’s correlation between the population response vectors of the two odors. The population response vector had length *l*, where *l* was the number of glomeruli, boutons, or cells in each experiment. Each element of this vector was the trial-averaged response of each glomerulus/bouton/cell to the stimulus. Representational dissimilarity matrices (RDM) show the pairwise correlation distances for every pair of odors imaged and aligned in the same experiment (same MB) and were computed using only the set of cells, glomeruli, or boutons in each experiment that responded to one or more odor stimuli. For each stage of processing, we computed a mean RDM by averaging the odor x odor RDMs across individual flies.

##### Comparison of representational space across individual MBs

For each MB, we generated an odor-odor distance vector of the correlation distances for every odor pair measured in common across the flies being compared. To evaluate the similarity of MB odor representational space across flies, we computed the Spearman’s rank correlation between the odor-odor distance vectors for every pairwise combination of MBs. As a control, the cell x odor response matrices for each MB were shuffled, and the same series of calculations applied. For each iteration of the shuffle, odor responses were randomized by permuting the columns of the cell x odor response matrix for each MB. The Spearman’s correlation computed from 10,000 iterations of this shuffle procedure was used to construct a distribution of Spearman’s correlations that would arise from chance for each pair of MBs, to be compared against the observed correlation.

##### Cell clustering

KCs were grouped by spectral clustering carried out on KC odor tuning profiles using ‘sklearn.cluster.SpectralClustering’^62^. Trial-averaged odor response vectors for all KCs that respond to one or more odors were pooled together from individual flies to create a grand cell x odor response matrix. The number of rows in this matrix equaled the sum of the number of responsive KCs across all flies and the number of columns equaled the number of odors that were sampled in common across all flies. A KC x KC affinity matrix was computed by taking the radial-basis transform of the KC x KC correlation distance matrix. Odor response profiles for each cluster were calculated by averaging the responses of all cells assigned to that cluster from each fly.

#### Modeling KC responses

We adapted a dynamic spiking model of the *Drosophila* olfactory network^30^ (https://github.com/annkennedy/mushroomBody) that implements the functional and anatomical organization of the circuit. ORN input to the model was derived from a published experimental dataset of the firing rate responses of 23 fly ORs to a panel of 109 odors^32^. The model implements lateral inhibition in the AL with a divisive inhibition term that normalizes PN firing rates. The model captures the response dynamics of PN and KC firing rates to an odor pulse, where ORN input is estimated by simply convolving a step to the steady-state firing rate with the cell’s synaptic membrane filter. For our analyses in this study, we focused on the mean firing rate of each cell (PN or KC) averaged over the odor pulse.

The model was implemented under several different assumptions of PN-KC connectivity. In the ‘uniform’ model, each of 2000 KCs had a mean of six input glomeruli, with each input independently assigned to a glomerulus at random. Each glomerulus had an equal likelihood of being selected at every input. In the ‘hemidraw’ model, each KC in the model (1748 cells) corresponded to a KC in the hemibrain connectome dataset^63^ and was assigned its observed number of claws (between 1 and 12, mean of 5.36 claws). Each claw was then independently assigned to a glomerulus according to the frequency of PN boutons for that glomerulus in the hemibrain MB; thus, glomeruli varied in their likelihood of being drawn as an input to each claw. In the ‘hemibrain’ model, each KC in the model was directly assigned the set of glomerular inputs of its corresponding cell in the hemibrain connectome. However, since glomeruli in the model were limited to the 23 ORs available in the Hallem dataset, modeled ‘hemibrain’ KCs had only an average of ~4 inputs. To evaluate the impact of a reduced number of inputs, versions of the ‘uniform’ and ‘hemidraw’ models were run with the distribution of the number of claws per KC centered on 4. These models predicted very similar odor-odor relationships to the earlier versions (data not shown).

The response rate of KCs to odor depends on both KC spiking threshold and the strength of feedback inhibition to KCs from the anterior paired lateral (APL) GABAergic neuron. In all versions of the model, KC spiking thresholds were assumed to be uniform across KCs and were set to achieve an average KC response rate of 20% across odors. Global inhibition by the APL was modeled as divisive inhibition at KC presynaptic terminals, and APL-KC weights were adjusted to halve the mean response rate across odors to 10%. This procedure was motivated by experimental observations that silencing APL output roughly doubles KC response rates.^22^ The APL was assumed to uniformly receive equal input from, and send equal output to, all KCs. For modeling of KC responses to CO_2_ (a stimulus not in the Hallem dataset), an additional glomerulus (corresponding to glomerulus V) and odor (CO_2_) was added into the OR input matrix. To estimate an upper bound for KC response rates to CO_2_, the firing rate of glomerulus V to CO_2_ was set to the maximum ORN firing rate (300 Hz) and set to zero for all other odors^57^.

#### Odor properties

Chemical descriptors were computed for each odor using the software Mordred^41^. Clustering over features identified a reduced set of 570 descriptors that captured odor-odor relationships in the full set of ~1800 descriptors; our analysis used this reduced set of descriptors (Supplemental Table S3). Odor relationships in natural odor space were estimated from a large database of headspace volatile profiles of natural food sources compiled from published datasets from the food science literature (Volatile Compounds in Food, VCF16.9 database, BeWiDo BV). The database contains 5564 observations from thousands of references. We focused on 2190 observations for which volatile profiles were quantitatively described in standardized units that could be compared between sources. For our analysis, we filtered compounds to isolate those present in ten or more observations (775 odors) and used the log scale of odor concentration.

#### Multidimensional scaling of odors in natural source space

We used hyperbolic non-metric multidimensional scaling (H-MDS)^64^ to embed odors into a low-dimensional hyperbolic space based on correlation distance, which reconstructs original distances monotonically (with preserved rank-ordering). The minimum Bayesian information criteria (BIC) determines the best dimension of the embedding. A hyperbolic metric was then used to measure distances between odors within the low-dimensional embedding space. To check that the results generalize across different subsets of natural odor sources, we repeated the analysis for a separate, albeit smaller secondary dataset that was compiled from another set of natural source literature references (see Supplemental Table S4). We find that odor pair distances computed in the VCF and in the secondary natural source dataset were positively correlated (rho=0.85, p=1.5e-8). The correlation between datasets increased with increasing the minimum number of odor sources for which a monomolecular odorant was required to be present in calculating distances between odorants. This result indicates increased stability of distances between more common odorants, that are ubiquitously present in natural environment. Furthermore, in both datasets, the hyperbolic space provided the best low-dimensional description. The curvature and best fitting dimensionality (dim=3) were similar, with curvature = −5.1 for the VCF dataset and curvature = −4.1 for the secondary natural source dataset.

#### Hierarchical clustering of odor representations

Representational dissimilarity matrices (RDM) show the pairwise correlation distances between every pairwise combination of odors imaged and aligned in the same experiment. For each stage of processing, a mean RDM was computed by averaging the odor x odor RDMs across individual flies. Dendrograms describing the relationships among odors at each stage of processing were generated from the mean odor x odor RDM for each stage. Hierarchical clustering of odors was implemented with ‘sklearn.cluster.AgglomerativeClustering’, using Pearson’s correlation as the distance metric and average linkage criterion (minimizes the average of the distances between all observations of pairs of clusters). These odor x odor RDM matrices were treated as feature matrices with dimensions (samples, features) rather than distance matrices – each odor was a different sample/row, and that odor’s features were its distance to the odors.

#### Data inclusion criteria

The number of flies (observations, *n*) in which each odor was measured is in Supplemental Table S1. The number of observations of each odor-odor distance (odor pair measured in the same MB) is in Supplemental Table S2. All flies analyzed in this study satisfied the following criteria. First, any field-of-view drift and warping of structure could be fully corrected using posthoc image registration, as evaluated by the ‘crispness’ of time-averaged movies (individual nuclei distinct and separated). Second, stimulus-evoked responses were reliably observed over the course of the entire recording in a ‘bulk’ fluorescence signal extracted in each frame from a global ROI circumscribing the entire imaged structure. Third, trial-trial correlation distances for repeated presentations of the same odor stimulus were clearly more similar to one another than for presentations of different odor stimuli. Sample sizes were not predetermined using a power analysis. We used sample sizes comparable to those used in similar types of studies^2,3^.

## ACKNOWLEDGEMENTS

We thank D. Anderson for sharing the *UAS-OpGCam6f* and *nls-OpGCaMP6s* fly lines. We thank A. Kennedy for modeling code and assistance with adapting the olfactory network model in this study. We thank V. Hauser for contributions to compiling natural odor source datasets. We thank P. Kandimalla for advising on connectomic analyses. We thank members of the Hong lab for careful readings of the manuscript. This work was supported by NIH grant R01MH117825 to E.J.H., and by the NSF/CIHR/DFG/FRQ/UKRI-MRC Next Generation Networks for Neuroscience Program (Award #2014217) to T.O.S. and E.J.H.

## AUTHOR CONTRIBUTIONS

J.Y.Y., T. F.O., T.O.S., and E.J.H. conceived the project, analyzed data, and wrote the manuscript. J.Y.Y. and K.V.D. performed imaging experiments in PN axons and KC somata. T.F.O. performed imaging experiments in ORN axons and PN dendrites and supervised curation of the natural odor source database. J.Y.Y., T.F.O., M.S.B., and W.M.H. analyzed data and generated figures. E.J.H. and T.O.S. supervised the project and acquired funding.

## DECLARATION OF INTERESTS

The authors declare no competing interests.

## SUPPLEMENTAL MATERIALS

**Supplementary Figure S1:**
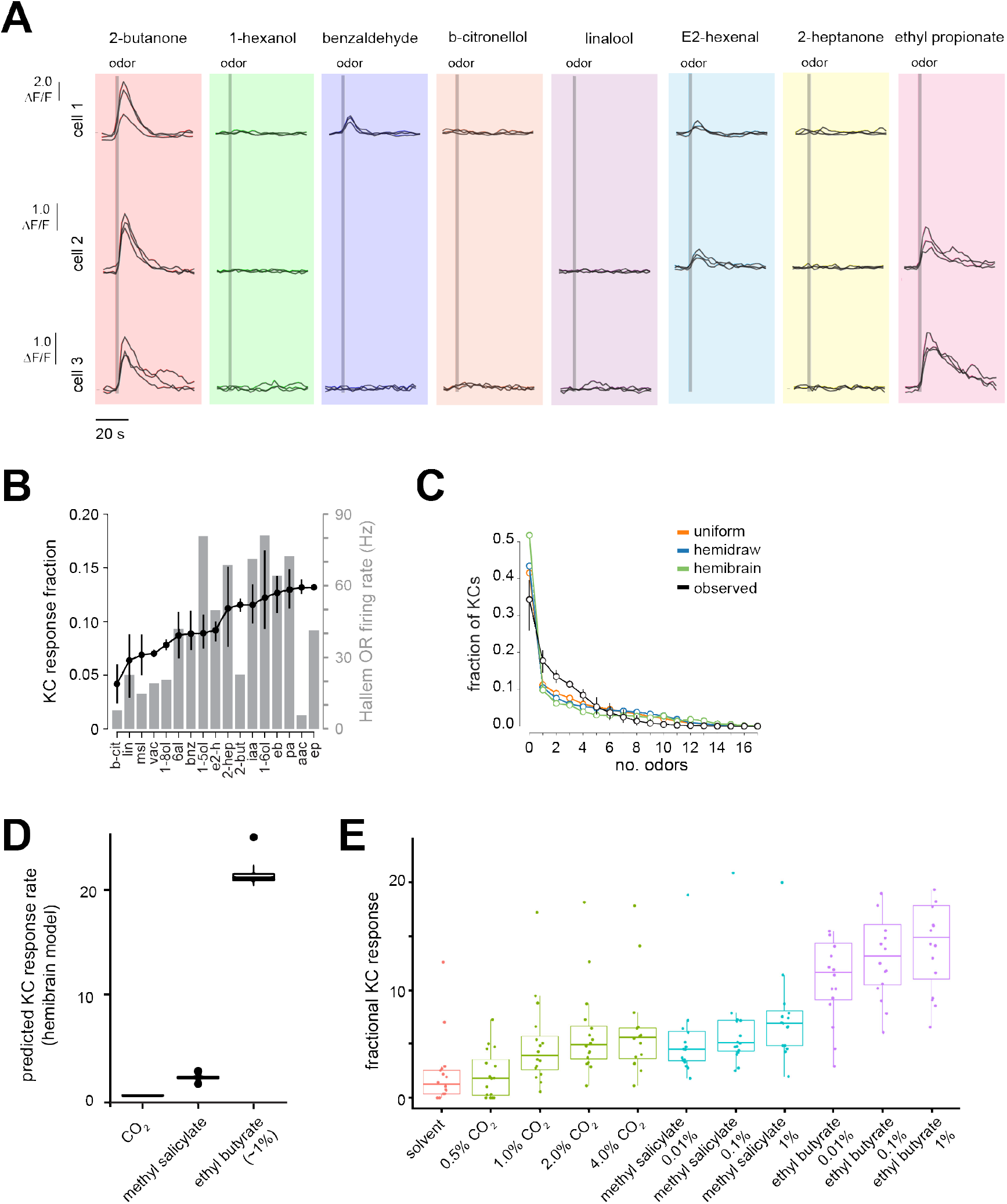
KC response properties. **A)** Odor-evoked responses in three example KCs from odor trials in different movies (imaging sessions with overlapping but distinct panels) collected in the same MB. Tracking and assignment of odor responses to KCs is reliable across imaging sessions. **B)** Comparison of KC response rate (mean and 95% CI, *n*=4 flies) and mean evoked OR firing rate (across glomeruli) in the Hallem dataset to each odor. **C)** Observed fraction of KCs (mean and 95% CI, *n*=4 flies) responding to the indicated number of odors for a panel containing 17 odors. The distribution of the fraction of modeled KCs responding to different numbers of odors in the uniform, hemidraw, and hemibrain models (see Figure 3) are plotted for comparison. **D)** Predicted KC response rates to narrowly activating (CO_2_, methyl salicylate) and broadly activating (ethyl butyrate) odors, under assumptions of uniform connectivity. For selective odors, the input firing rate for the cognate OR was set to saturating firing rates (300 Hz) to estimate the upper bound for KC response rate. For ethyl butyrate, the observed firing rates across ORs in the Hallem dataset was used as input to the model. **E)** Observed KC response rates to varying concentrations of the odors in **D**.

**Figure S2:**
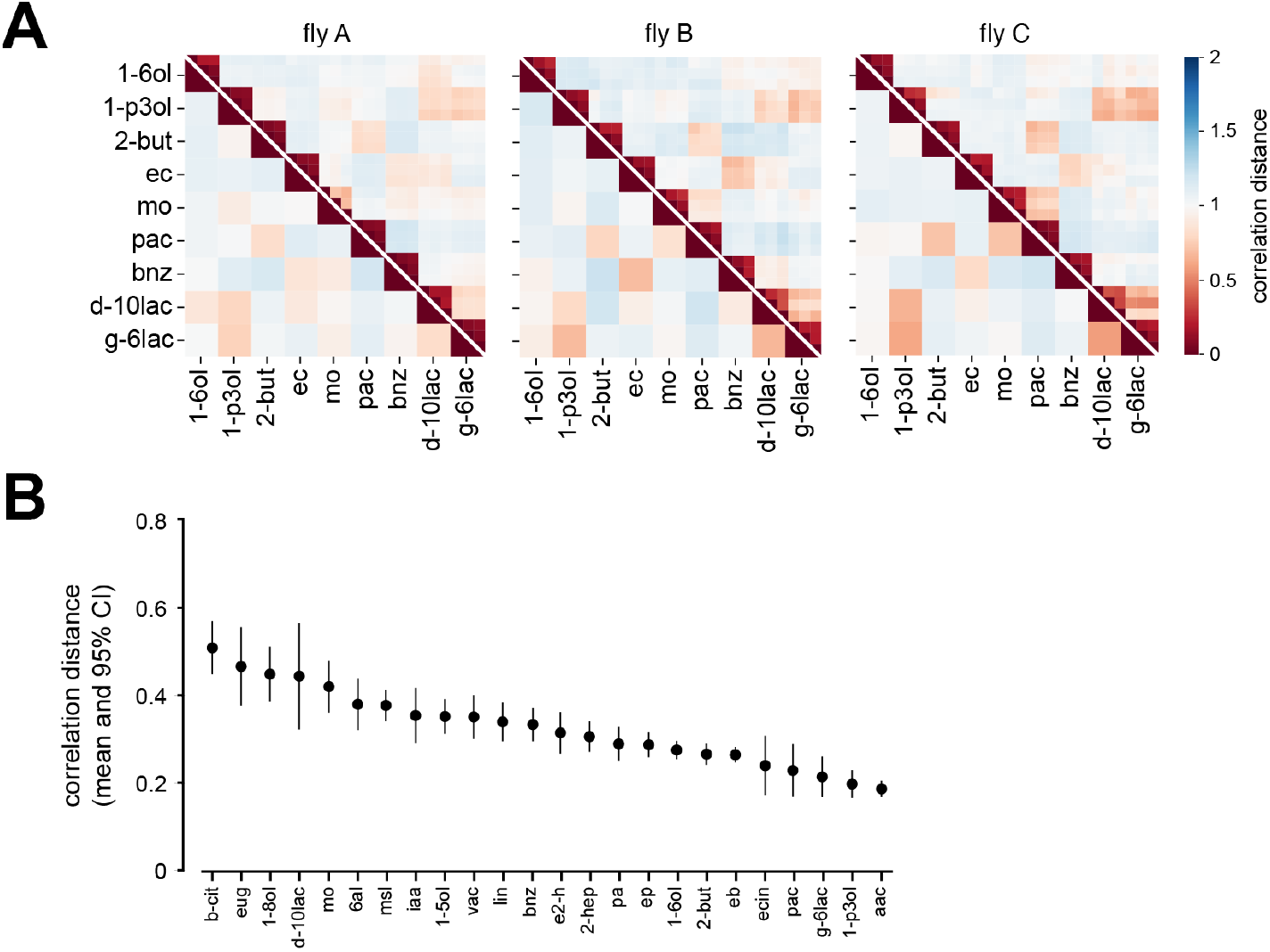
KC population responses to another example odor panel, illustrating the invariance of MB representational space across individuals. **A)** Matrices of correlation distances from three different flies showing pairwise relationships between KC population responses in individual odor trials (upper triangles) or in trial-averaged responses for each odor (lower triangle). **B)** Correlation distances between KC response vectors from different trials of the same odor. For each odor, the trial-trial KC response correlation was computed for all pairs of trials of this odor in each fly and averaged. The plot shows the mean and 95% CI of the fly averages. Compare to Figure 2E. Weaker odors tend to be less reliable than stronger odors.

**Figure S3:**
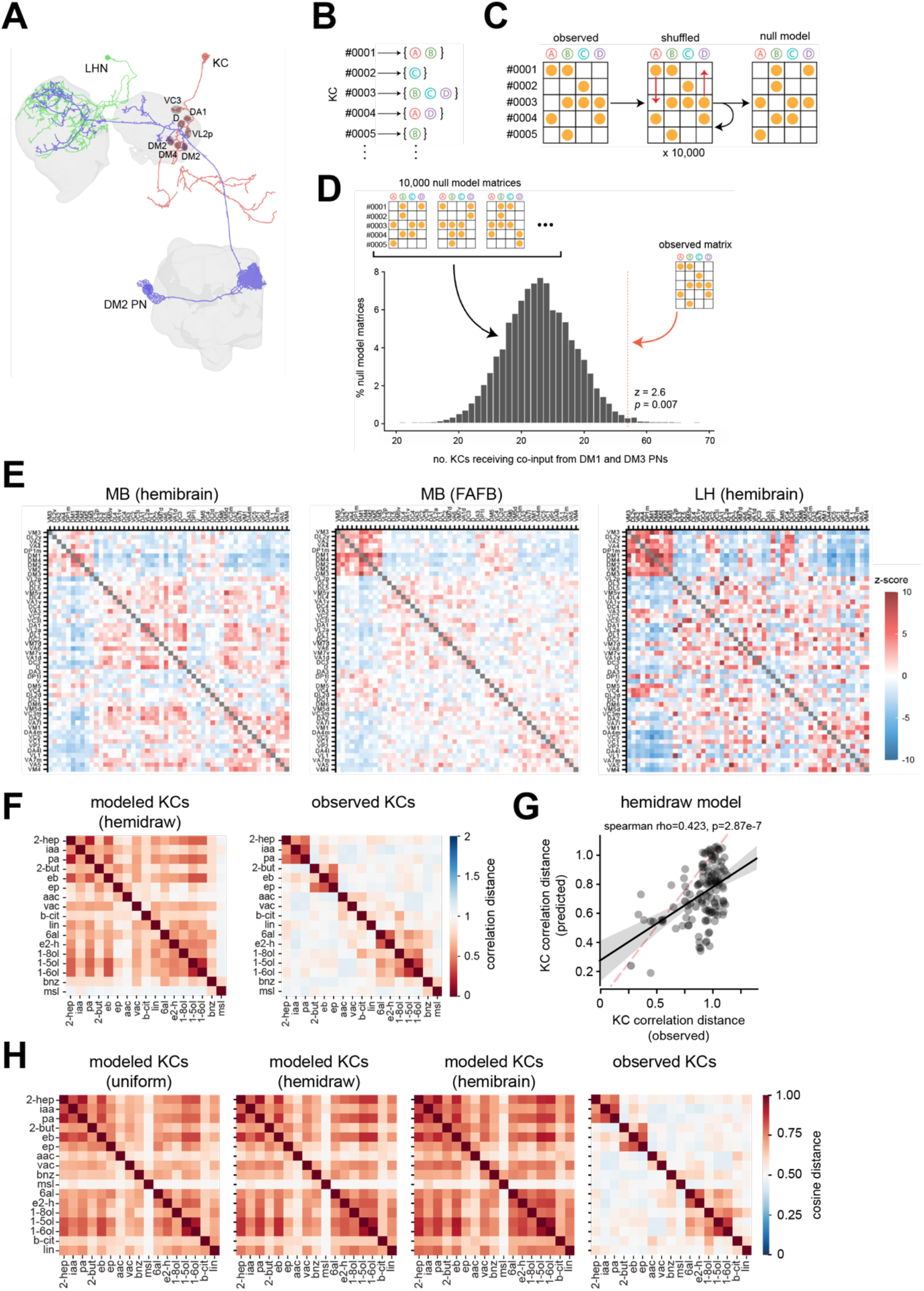
The structure of olfactory glomerular inputs to third-order olfactory neurons and the impact on MB representations of odor. **A)** Example reconstruction from the hemibrain connectome^63^ of a uniglomerular DM2 PN and two example postsynaptic partners, a Kenyon cell (KC) and a lateral horn neuron (LHN). Synaptic connections between identified neurons contained within the hemibrain volume are fully described. **B)** For each third-order olfactory neuron, the set of glomeruli providing direct presynaptic input via PNs was extracted from connectome datasets (hemibrain and FAFB). For this analysis, synaptic connections were binarized (e.g., all glomerular inputs >5 synapses were treated equally regardless of synapse count). **C)** The mapping of glomerular inputs to KCs was represented as a binary matrix, where a 1 in cell(*i,j*) indicates that neuron *i* receives input from glomerulus *j*. The ‘Curveball’ algorithm was used to generate random matrices (null model) that preserve the row and column totals of the original matrix^65^. **D)** For each pair of glomeruli, a distribution of the number of KCs receiving co-input from the two glomeruli in each shuffled matrix was generated. The observed number of KCs receiving co-input from each pair of glomeruli in the hemibrain was compared to this distribution to generate a z-score. An example distribution and z-score for the over-convergent glomerular pair DM1 and DM3 are given. **E)** Glomerular input structure to KCs in the hemibrain MB^63^ (left), KCs in the FAFB MB^66^ (center), and lateral horn neurons in the hemibrain LH (right). The z-score for each pair of glomeruli measures the degree to which the glomeruli are over- or under-convergent in the observed population, compared to null models. A large positive value indicates strong over-convergence, and a large negative value indicates strong under-convergence. The ordering of glomeruli is the same in all matrices and was based on k-means clustering of the hemibrain MB matrix. **F)** Matrix of pairwise correlation distances between predicted KC responses to 17 odors in the hemidraw model (left). The mean pairwise relationships for observed KC responses are reproduced from Figure **3Ciii** (right) for ease of comparison. **G)** Comparison of odor-odor correlation distances between observed KC responses and predicted KC responses in the hemidraw model. Each symbol is an odor pair. **H)** Matrix of pairwise cosine distances between predicted KC responses (uniform, hemidraw, or hemibrain models), or observed KC responses, for 17 odors.

**Figure S4:**
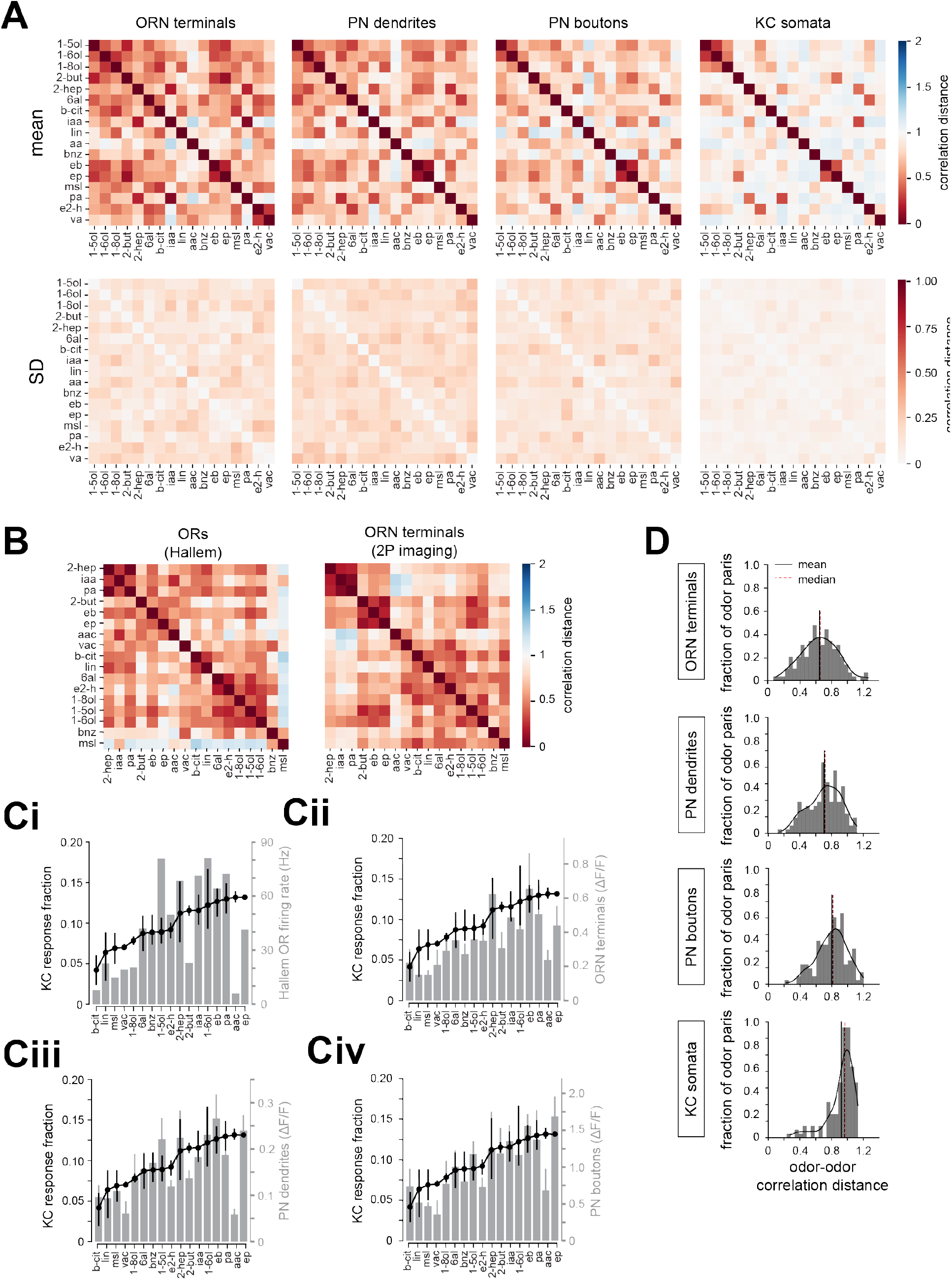
Transformation of odor representations in the fly olfactory circuit. **A)** Mean (top row) and standard deviation (bottom row) across flies of pairwise correlation distances of odor representations in ORN axons (*n* = 4 flies), PN dendrites (*n* = 6 flies), PN boutons (*n* = 6 flies), and KC somata (*n* = 4 flies) for a panel of 17 odors. **B)** Pairwise correlation distances of OR tuning in the Hallem dataset (left) and ORN axon terminal responses measured in this study (right) for a panel of 17 odors. **C)** Comparison of fractional KC response rates with odor-evoked **Ci**) firing rates across ORs in the Hallem dataset (grey); **Cii**) ORN terminal responses (grey); **Ciii**) PN dendrite responses (grey); and **Civ**) PN bouton responses (grey) for 17 odors. ORN terminal, PN dendrite, PN bouton, and KC responses are mean and 95% CI across flies of the ROI-averaged evoked response to each odor in each fly. **D)** Distribution of odor-odor correlation distances for representations at each stage of olfactory processing.

**Supplemental Table S1:** Odor information and number of observations (flies) for each odor in KC somata datasets.

| Odor | Abbreviation | InChI | Sigma-Aldrich Cat. No. | CAS | No. flies sampled (n) |
| --- | --- | --- | --- | --- | --- |
| 1-hexanol | 1-6ol | InChI=1S/C6H14O/c1-2-3-4-5-6-7/h7H,2-6H2,1H3 | 471402 | 111-27-3 | 22 |
| 1-octanol | 1-8ol | InChI=1S/C8H18O/c1-2-3-4-5-6-7-8-9/h9H,2-8H2,1H3 | 297887 | 111-87-5 | 16 |
| 1-pentanol | 1-5ol | InChI=1S/C5H12O/c1-2-3-4-5-6/h6H,2-5H2,1H3 | 138975 | 71-41-0 | 10 |
| 1-penten-3-ol | 1-p3ol | InChI=1S/C5H10O/c1-3-5(6)4-2/h3,5-6H,1,4H2,2H3 | 1984 | 616-25-1 | 7 |
| 2-butanone | 2-but | InChI=1S/C4H8O/c1-3-4(2)5/h3H2,1-2H3 | 360473 | 78-93-3 | 21 |
| 2-heptanone | 2-hep | InChI=1S/C7H14O/c1-3-4-5-6-7(2)8/h3-6H2,1-2H3 | 537683 | 110-43-0 | 14 |
| acetic acid | aac | InChI=1S/C2H4O2/c1-2(3)4/h1H3,(H,3,4) | 695092 | 64-19-7 | 5 |
| B-citronellol | b-cit | InChI=1S/C10H20O/c1-9(2)5-4-6-10(3)7-8-11/h5,10-11H,4,6-8H2,1-3H3 | C83201 | 106-22-9 | 12 |
| benzaldehyde | bnz | InChI=1S/C7H6O/c8-6-7-4-2-1-3-5-7/h1-6H | 418099 | 100-52-7 | 20 |
| delta-decalactone | d-10lac | InChI=1S/C10H18O2/c1-2-3-4-6-9-7-5-8-10(11)12-9/h9H,2-8H2,1H3 | 74026 | 705-86-2 | 5 |
| E2-hexenal | e2-h | InChI=1S/C6H10O/c1-2-3-4-5-6-7/h4-6H,2-3H2,1H3/b5-4+ | 158131000 | 6728-26-3 | 11 |
| ethyl butyrate | eb | InChI=1S/C6H12O2/c1-3-5-6(7)8-4-2/h3-5H2,1-2H3 | E15701 | 105-54-4 | 14 |
| ethyl cinnamate | ec | InChI=1S/C11H12O2/c1-2-13-11(12)9-8-10-6-4-3-5-7-10/h3-9H,2H2,1H3/b9-8+ | 66761 | 103-36-6 | 5 |
| ethyl propionate | ep | InChI=1S/C5H10O2/c1-3-5(6)7-4-2/h3-4H2,1-2H3 | 112305 | 105-37-3 | 16 |
| eugenol | eug | InChI=1S/C10H12O2/c1-3-4-8-5-6-9(11)10(7-8)12-2/h3,5-7,11H,1,4H2,2H3 | E51791 | 97-53-0 | 7 |
| gamma-hexalactone | g-6lac | InChI=1S/C6H10O2/c1-2-5-3-4-6(7)8-5/h5H,2-4H2,1H3 | 68554 | 695-06-7 | 6 |
| hexanal | 6al | InChI=1S/C6H12O/c1-2-3-4-5-6-7/h6H,2-5H2,1H3 | 115606 | 66-25-1 | 15 |
| isoamyl acetate | iaa | InChI=1S/C7H14O2/c1-6(2)4-5-9-7(3)8/h6H,4-5H2,1-3H3 | W205532 | 123-92-2 | 12 |
| linalool | lin | InChI=1S/C10H18O/c1-5-10(4,11)8-6-7-9(2)3/h5,7,11H,1,6,8H2,2-4H3 | L2602 | 78-70-6 | 10 |
| methyl octanoate | mo | InChI=1S/C9H18O2/c1-3-4-5-6-7-8-9(10)11-2/h3-8H2,1-2H3 | 21719 | 111-11-5 | 9 |
| methyl salicylate | msl | InChI=1S/C8H8O3/c1-11-8(10)6-4-2-3-5-7(6)9/h2-5,9H,1H3 | M6752 | 119-36-8 | 14 |
| pentyl acetate | pa | InChI=1S/C7H14O2/c1-3-4-5-6-9-7(2)8/h3-6H2,1-2H3 | 109584 | 628-63-7 | 15 |
| propyl acetate | pac | InChI=1S/C5H10O2/c1-3-4-7-5(2)6/h3-4H2,1-2H3 | 40858 | 109-60-4 | 7 |
| valeric acid | va | InChI=1S/C5H10O2/c1-2-3-4-5(6)7/h2-4H2,1H3,(H,6,7) | 240370 | 109-52-4 | 13 |

**Supplemental Table S2:** Mean, 95% CI, SEM, and number of observations (flies) of the correlation distance for every unique odor pair in KC somata datasets.

**Supplemental Table S3:** Reduced list of Mordred molecular descriptors used in this study.

**Supplemental Table S4:** References contributing to a secondary natural odor source database.

## Notes

### Competing Interest Statement

The authors have declared no competing interest.

### Summary of Updates

The revision corrects minor formatting and textual errors in the first version.

